# Dissecting the molecular basis of variability for flowering time in *Camelina sativa*

**DOI:** 10.1101/2024.11.05.620108

**Authors:** Liyong Zhang, Venkatesh Bollina, Peng Gao, Isobel A. P. Parkin

**Affiliations:** Agriculture and Agri-Food Canada, 107 Science Place, Saskatoon, SK S7N0X2, Canada

**Keywords:** Camelina, flowering time, pan-transcriptome, genetic variation, association analyses

## Abstract

*Camelina sativa* is an important polyploid oilseed crop with multiple favorable agronomic traits. Capturing the leaf transcriptome of 48 accessions of *C. sativa* suggests allelic variation for gene expression levels and notably sub-genome dominance, both of which could provide opportunities for crop improvement. Flowering time (FT) is a crucial factor affecting the overall yield of crops. However, our understanding of the molecular mechanisms underlying FT regulation in *C. sativa* are still limited, partly due to its complex allohexaploid genome. In this study, weighted gene co-expression network analysis (WGCNA), expression quantitative trait loci (eQTL) analysis and transcriptome-wide association study (TWAS) were employed to explore the FT diversity among 48 *C. sativa* accessions and dissect the underlying molecular basis. Our results revealed a FT-related co-expressed gene module highly enriched with *SOC1*s and *SOC1*-like genes, and identified 10 significant FT-associated single nucleotide polymorphisms (SNPs) defining three haplotype groups; thus providing a molecular basis for future genetic improvements in *C. sativa* breeding.

## Introduction

Like the model plant *Arabidopsis thaliana, Camelina sativa* is a member of the Brassicaceae family. Due to the sustainable and adaptable nature of the crop, over the past decade *C. sativa* has gained considerable attention as an important oilseed to support the food, feed and biofuel industries (Berti et al., 2016; Vollmann & Eynck, 2015). The genome sequence of *C. sativa* was published in 2014 (Kagale et al., 2014) and a transcriptome atlas detailing gene expression across a range of tissues has been developed (Kagale et al., 2016), providing a knowledge base for further development of the crop. Yet trait improvement relies on access to underlying genetic variation and a number of genotyping studies have shown limited variation within available *C. sativa* germplasm (Chaudhary et al., 2020; Gehringer et al., 2006; Luo et al., 2019; Singh et al., 2015). A recent assessment of genetic diversity in the largest collection of lines to date concluded that *C. sativa* has a modest degree of genome diversity, comparable to that of related *Brassica napus* (Li et al., 2021). This would concur with the evolutionary trajectory of the two crops, both being recent neopolyploids, probably formed from a limited number of hybridization events. Despite the modest level of genome diversity, Li et al (2021) identified a wide range of phenotypic variation for a number of important agronomic traits. The phenotypic variation could stem from the complex control of many traits by a large number of small effect loci, which was suggested by the inherent difficulty in applying genome wide association analyses (GWAS) in *C. sativa* (Li et al., 2021).

At the level of gene expression there is limited information available for variation among *C. sativa* accessions with data collected from only a handful of lines (Gomez-Cano et al., 2020). Many lines of evidence implicate gene regulation as the source of variation for a large number of traits and the application of GWAS based on transcriptome data has identified useful loci controlling traits in species with recognized low genetic diversity, such as *B. napus* (Harper et al., 2012; Havlickova et al., 2018; Tang et al., 2021). In the current study, transcriptome data was collected for 48 publicly available accessions of *C. sativa* in order to assess the level of available variation and to provide an exploitable resource for researchers.

For plants, the transition from vegetative to reproductive phase, or flowering, is one of the most important developmental switches from the perspective of reproductive success. Regulation of flowering has been investigated over many decades in the model plant *Arabidopsis thaliana* and other flowering plants. Tremendous progress have been made with six different flowering pathways established, namely; autonomous, vernalization, photoperiod, aging, gibberellin (GA), and thermosensory pathways. In addition, there are complex interactions between the pathways, which determine reproductive success. For instance, both the autonomous and thermosensory pathways promote the floral transition through the main floral repressor, FLOWERING LOCUS C (FLC) (Cheng et al., 2017; Simpson 2004; Capovilla et al., 2014; Michaels and Amasino, 1999; Sheldon et al., 1999). Increasing evidence has shown these floral pathways converge on several common genes, well-known as floral integrators, including *FLOWERING LOCUS C* (*FLC*), *CONSTANS* (*CO*), *SUPPRESSOR OF OVER EXPRESSION OF CONSTANS* (*SOC1*) and *LEAFY* (*LFY*) to fine-tune flowering time (Moon 2005; Araki 2001; Komeda 2004; Srikanth and Schmid 2011). In addition, epigenetic pathways and small non-coding RNAs have also been shown to play important roles in flowering time regulation (Wang and Köhler 2017; Teotia and Tang 2015). Thus, flowering plants have developed an elaborate network of genetic pathways responding to both exogenous environmental stimuli (e.g. temperature and photoperiod) and endogenous cues (e.g. hormone, sugars) to regulate floral transition.

However, for the allohexaploid *C. sativa*, a close relative of *A. thaliana*, there have been limited studies of how flowering time is regulated. Given that the floral integrator genes (e.g. *FT* and *SOC1*) are highly conserved among most of the flowering plants, and the high collinearity between *C. sativa* and *A. thaliana* genomes (Kagale et al., 2014), researchers have attempted to understand the molecular basis underlying flowering time in *C. sativa* through isolating homologues of key flowering time genes. Leveraging available genetic and genomic tools, a number of studies have identified the orthologues of the key floral repressor gene *FLOWERING LOCUS C* (*FLC*) in *C. sativa* as playing an important role in the flowering habits of *C. sativa* (Anderson et al., 2018; Chao et al., 2019; Chaudhary et al., 2023). Luo Lily et al. (2021), reported 20 significant flowering time-associated single nucleotide polymorphisms (SNPs) through a genome-wide association study (GWAS) alone (Luo Lily et al., 2021). As the natural diversity of *C. sativa* provides an important revenue for positive trait selection, and large variation in flowering time have been documented among different *C. sativa* accessions (Berti et al., 2016; Chao et al., 2019; Luo Lily et al., 2021); in this study, we exploited the developed transcriptome resource from the 48 different genotypes by integrating WGCNA, eQTL and TWAS analysis to dissect the molecular basis underlying flowering time regulation among these lines. Our results provide insights into the regulatory mechanisms underlying *C. sativa* FT trait and present 10 FT-associated SNPs, which could provide a basis for modifying FT during future *C. sativa* breeding.

## Results

### 1 Gene expression analyses among *C. sativa* accessions

To facilitate the investigation of the role of gene regulation in controlling expression of agronomic traits RNA sequencing was conducted with young leaf tissue from 48 genotypes. Overall, approximately 926 million clean reads were obtained, with an average number of 18.9 million reads per genotype. All clean reads were aligned against the *C. sativa* reference genome, with mapping rates ranging from 72.5% to 82.9%. After quantifying the gene expression levels, of the total 94,495 *C. sativa* gene models, 37,268 were detected across all 48 genotypes and 72,713 genes were expressed in at least at one genotype, while the numbers of expressed genes varied between different genotypes ranging from 45,642 (CN113724) to 55,582 (CN113692) (**Supplementary Figure S1A**). The gene expression data provides an indication of the level of variability in the population. Treating the data as a pan-transcriptome dataset identified a subset of 184 highly variable genes that are significantly dominantly (or specifically) expressed in 10 or fewer of the lines (TAU>0.9) (**Figure 1**). GO annotation enrichment of this gene set showed a significant over-representation of genes involved in stress responses, in particular in plant defense (**Supplementary Figure S2**). Interestingly some lines show expression of a high proportion of these defense genes.

**Figure 1.**
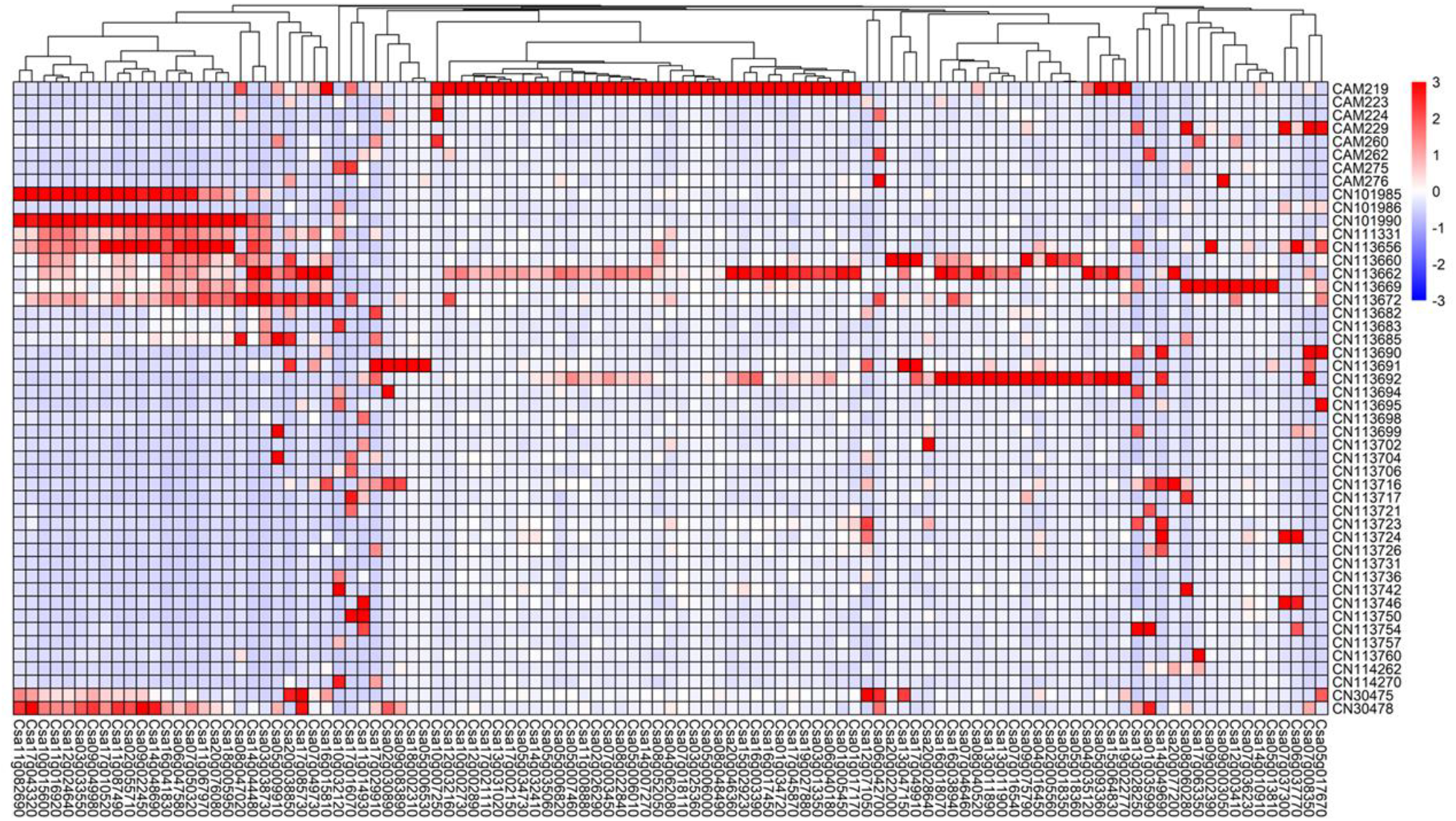
Dominantly expressed genes across the 48 *C. sativa* accessions. 184 highly variable genes (x-axis) identified based on TAU index were Z-score normalized and then visualized using hierarchical clustering combined with a gene expression heatmap. The color scheme, from red through white to blue, indicates the level of normalized expression, from high to low across 48 *C. sativa* accessions (y-axis).

The polyploid nature of the *C. sativa* could provide another mechanism for creating expression diversity, since differential sub-genome bias or genome dominance is a common trait in polyploids and has previously been reported for the *C. sativa* reference genome (Kagale et al., 2014). Considering the 12,440 syntenic triads, consisting of orthologous genes from the three sub-genomes with some level of gene expression, 2,101 showed balanced genome expression across the 48 accessions and 2,460 showed the same pattern of genome dominance across the 48 lines (**Figure 2A**). The remaining triads showed differential patterns of genome dominance biased to one of the three sub-genomes among the 48 accessions (**Figure 2B**). For those triads where the bias was not conserved among the accessions, 284 showed significantly divergent expression patterns with less than half of the accessions displaying the same pattern of dominance (**Figure 2C**). Annotation of these divergent genes identified a wide range of transcription factors, potentially impacting a number of traits including flowering time (**Figure 2C**).

**Figure 2.**
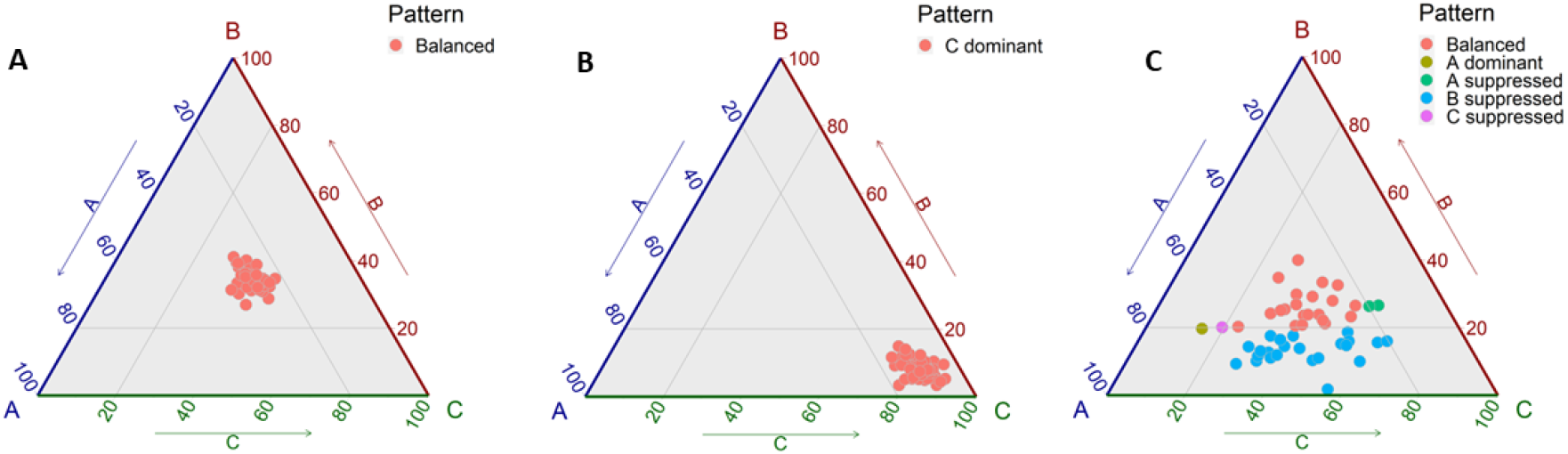
Patterns of Genome Dominance in 48 *C. sativa* accessions. The vertices of the triangle, A, B and C, represent the three orthologous gene copies in the sub-genomes of *C. sativa* respectively. Each filled circle represents the relative level of expression for each of the three genes and are coloured according to the shown legend. (A) Balanced gene expression for Csa11g001340, Csa10g001220 and Csa12g001280; (B) Sub-genome 3 genome dominance for triad Csa11g101280, Csa18g037800 and Csa02g072860; and (C) Divergent genome dominance among accessions for triad Csa14g063510, Csa03g060290 and Csa17g093410; orthologues of At1g53160 or SPL4, a FT gene.

### 2 Generation of genome-wide gene-associated SNPs

To generate genome-wide gene-associated single nucleotide polymorphisms (SNPs), all clean reads of above RNA sequencing were aligned against the *C. sativa* reference genome to generate bam files, which were used for calling SNPs through genome analysis toolkit (GATK). In total, 1,151,672 SNPs were identified, which were further filtered to 65,082 SNPs. Among the 65,082 SNPs, the majority (56,469) were located within exons, and 6,865 SNPs were in 3^’^ or 5^’^ untranslated regions, while introns contained the remaining 1,748 SNPs (**Supplementary Figure 1B**). The final 65,082 SNPs were used to perform a phylogenetic analysis which resolved the 48 genotypes of the experimental population into 4 distinct groups (**Figure 3A**).

**Figure 3.**
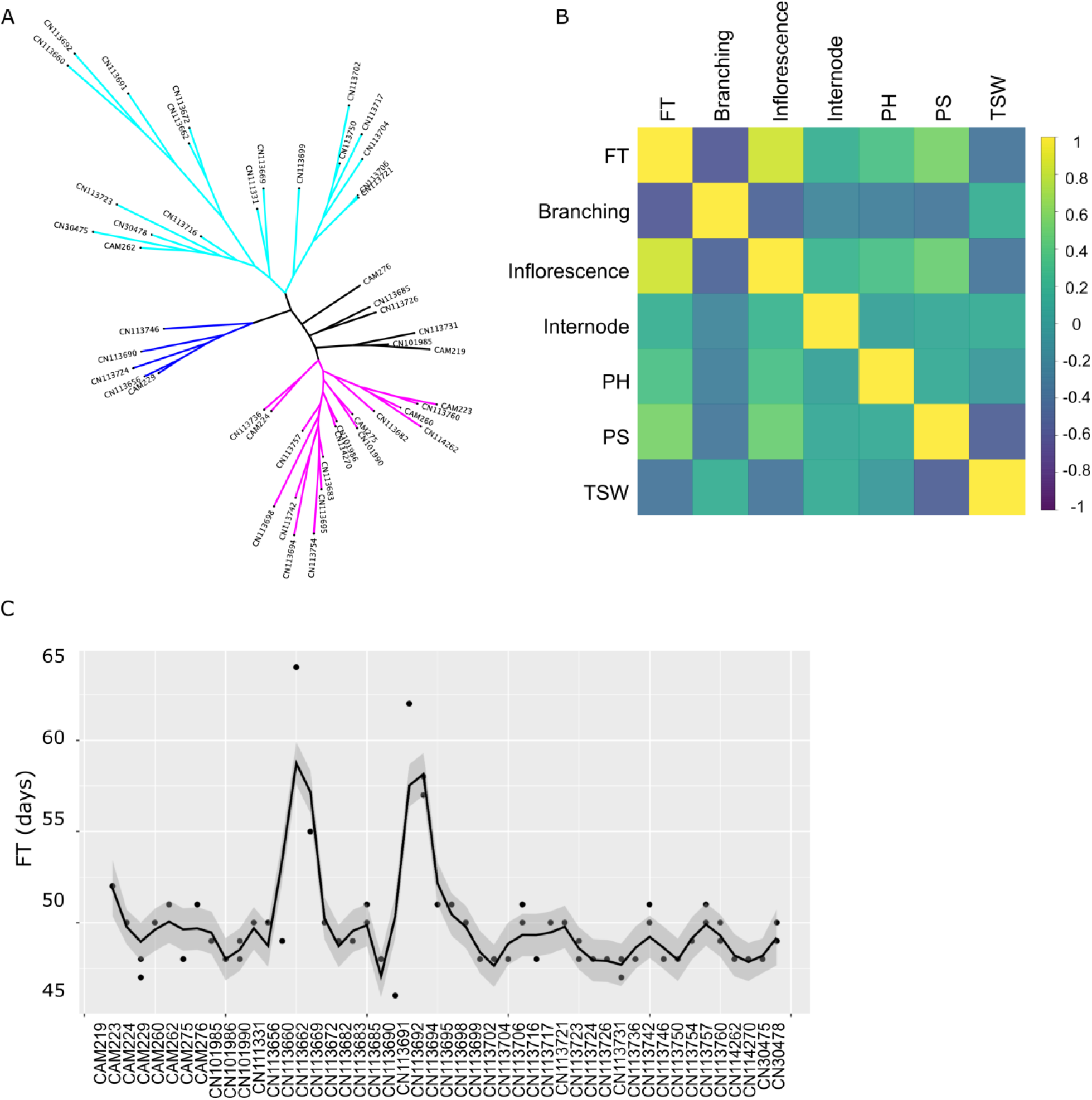
*C. sativa* population structure and FT variation. (A) Phylogenetic tree of all 48 experimental genotypes according to the final 65,082 SNPs. (B) Heatmap showing Pearson’s correlation between different traits. Flowering time, plant height, pod shattering, and 1000-seed weight represented by the abbreviation FT, PH, PS, and TSW respectively. (C) Smoothed line plot of FT across all 48 genotypes in 2012.

### 3 Trait variation across *C. sativa* population

The 48 accessions were grown in the field for two consecutive seasons in 2012 and 2013, and data for a number of traits was collected, including flowering time (FT), inflorescence and internode length, plant height, 1000-seed weight, pod shattering and branching (**Supplementary data 1**). The pairwise correlations between different traits were calculated using their best linear unbiased prediction values. As shown in **Figure 3B**, there was a significant positive correlation between FT and inflorescence length with a correlation coefficient of 0.84 (P-value < 3.4 × 10^−14^), suggesting these two traits are likely mutually dependent. Meanwhile, we also noticed a significant negative correlation between FT and branching with a correlation coefficient of -0.57 (P-value < 2.2 × 10^−5^)(**Figure 3B**). In the current study we have focused on FT during *C. sativa* development. To check FT variation across the two seasons, their correlation was calculated as Pearson’s correlation coefficient (*r* = 0.94 and p-value < 2.2 × 10^−16^), suggesting a high consistency for FT between the two environments in 2012 and 2013. For clarity, only data from year 2012 was shown in **Figure 3C**, whereat FT varies largely between different genotypes of *C. sativa*, e.g. CN113690 only needs 46 days to reach 50 % flowering on average, while CN113660 needs 8 more days (**Figure 3C**; see same trend of 2013 in **Supplementary Figure 1C**).

### 4 Identification of FT-related highly co-expressed gene modules

Weighted gene co-expression network analysis (WGCNA) provides a useful method to construct a co-expression network, where each module of the network contains a group of highly correlated genes, which tend to be involved in similar biological processes (Langfelder & Horvath, 2008). To explore the potential co-expressed gene module responsible for the above FT variation, we employed WGCNA to analyze the 15,000 genes with the highest expression variance across the experimental population. In the co-expressed network, 15,000 genes were divided into 19 distinct modules (**Figure 4A**) with a size range from 46 to 4,194 genes (**Table 1**). To quantify whether these modules associate with the FT trait, the eigengenes (i.e. those genes representative of gene expression profiles in a module) of all 19 modules were used to calculate their Pearson’s correlation coefficients with the FT trait (**Figure 4B**). As seen in **Figure 4B**, FT is highly negatively correlated with the tan module, which contains 164 *C. sativa* genes (**Table 1**). To further explore the genes in the tan module, the definitions from (Zhang & Horvath, 2005) were adapted, whereat the absolute value of the Pearson correlation between each gene and FT was defined as gene significance (GS), and module membership (MM) was defined by the individual gene’s correlation with the eigengene of the tan module. By quantifying the GS and MM for each gene in the tan module and plotting their GS against MM, it was clear that there was a strong positive correlation between GS and MM, which indicated that genes related to FT were also the central genes of the tan module (**Figure 4C**).

**Table 1:** Co-expressed gene modules and their gene numbers resulting from WGCNA.

| *Module* | grey | turquoise | blue | brown | yellow | green | red | black | pink | magenta |
| --- | --- | --- | --- | --- | --- | --- | --- | --- | --- | --- |
| *Number of genes* | 4194 | 3289 | 1133 | 1005 | 772 | 738 | 663 | 654 | 572 | 492 |
|  | purple | greenyellow | tan | salmon | cyan | midnightblue | lightcyan | grey60 | lightgreen |  |
|  | 355 | 341 | 164 | 160 | 140 | 115 | 87 | 80 | 46 |  |

**Figure 4.**
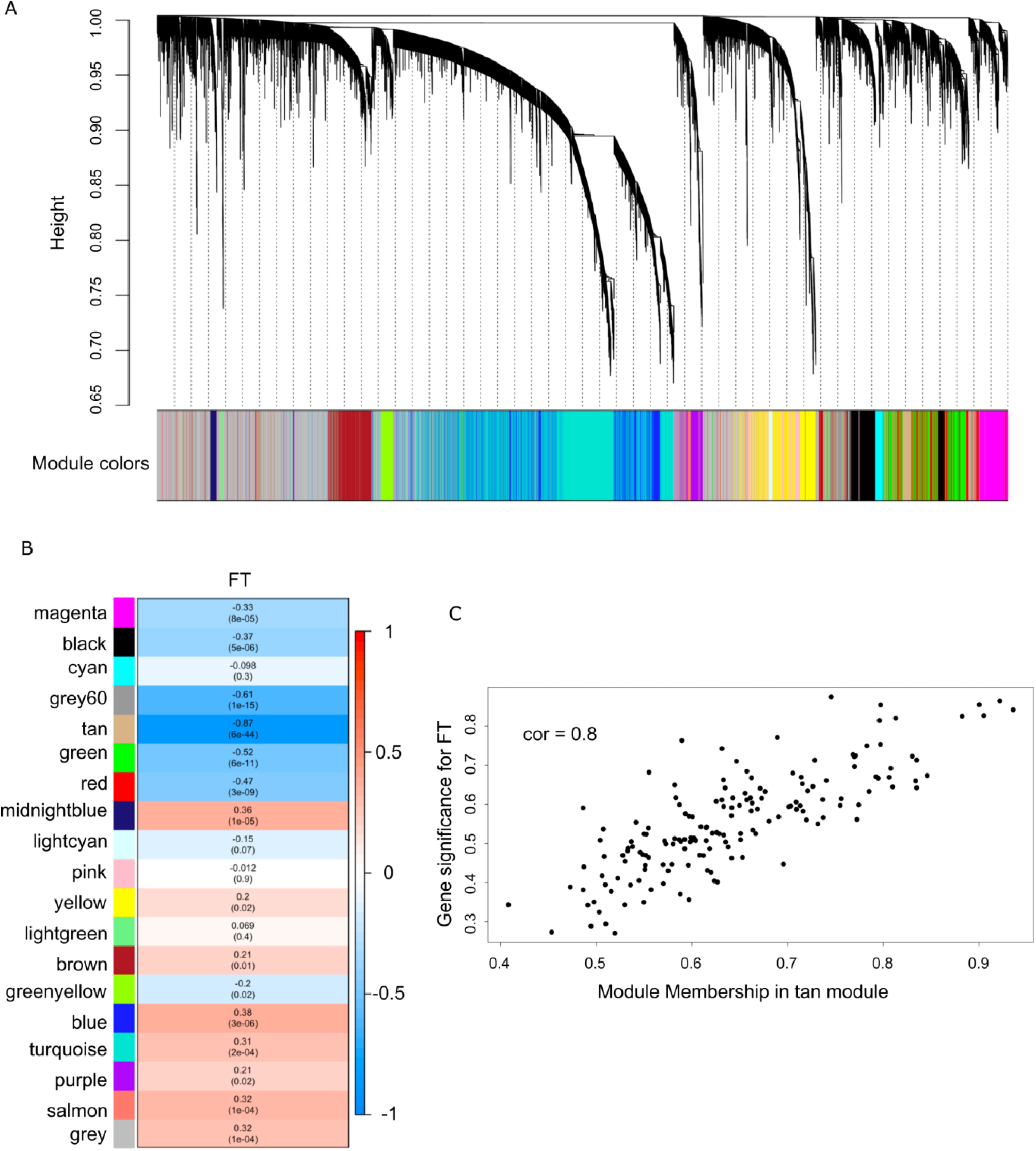
WGCNA identifies FT-related co-expressed gene module. (A) Hierarchical clustering dendrogram showing 19 distinct co-expression modules based on 15,000 most variable genes across 48 genotypes. Different colors were assigned to each module. (B) Heatmap showing the correlation of FT with each co-expression module. Each cell contains the corresponding Pearson’s correlation coefficient and p-value. (C) Scatterplot showing gene significance for FT and module membership in tan module.

After generating the gene ontology (GO) database of *C. sativa* genes (see Methods for details), GO enrichment analysis through clusterProfiler was used to functionally characterize the tan module. Five GO terms were significantly enriched in the tan module (P-values < 0.05 after false discovery rate correction) (**Figure 5A**). Listing all genes assigned to the enriched GO categories showed that, the categories “defense response to other organism” and “ADP binding” shared multiple genes, while the remaining enriched GO terms did not overlap (**Figure 5B**). After further examination, it was noticed that all genes assigned to the categories “RNA polymerase II transcription regulatory region sequence-specific DNA-binding” were members of the MADS transcription factor family, which are recognized for their role in floral organ development (Becker & Theissen, 2003). The MADS transcription factors enriched in the tan module are *AGL19* (AT4G22950) and *AGL20* (AT2G45660), both of which have been documented to participate in floral transition and promote flowering (Borner et al., 2000; Lee et al., 2000; Schonrock et al., 2006). Noticeably, *SOC1* (*AGL20*) is a well-known key regulator of flowering timing and integrates different flowering pathways, and *AGL19* is classified as *SOC1*-like gene according to its high similarity with *SOC1* (Becker & Theissen, 2003; Komeda, 2004). Perhaps not surprisingly due to the polyploid nature of *C. sativa*, the tan module contains more than one *SOC1*; Csa04g063650.1 and Csa06g052060.1 (**Figure 5B**; **Supplementary data 2**). More interestingly, the tan module also contains two *SOC1*-like genes, *AGL19*s (Csa10g020140.1 and Csa12g033740.1), which suggests these *C. sativa SOC1*s and *SOC1*-like genes might work closely together in a similar manner as their Arabidopsis orthologs. Thus, the network analysis identified a FT-related co-expressed gene module, which is highly enriched with MADS transcription factors, especially *SOC1*s and *SOC1*-like genes.

**Figure 5.**
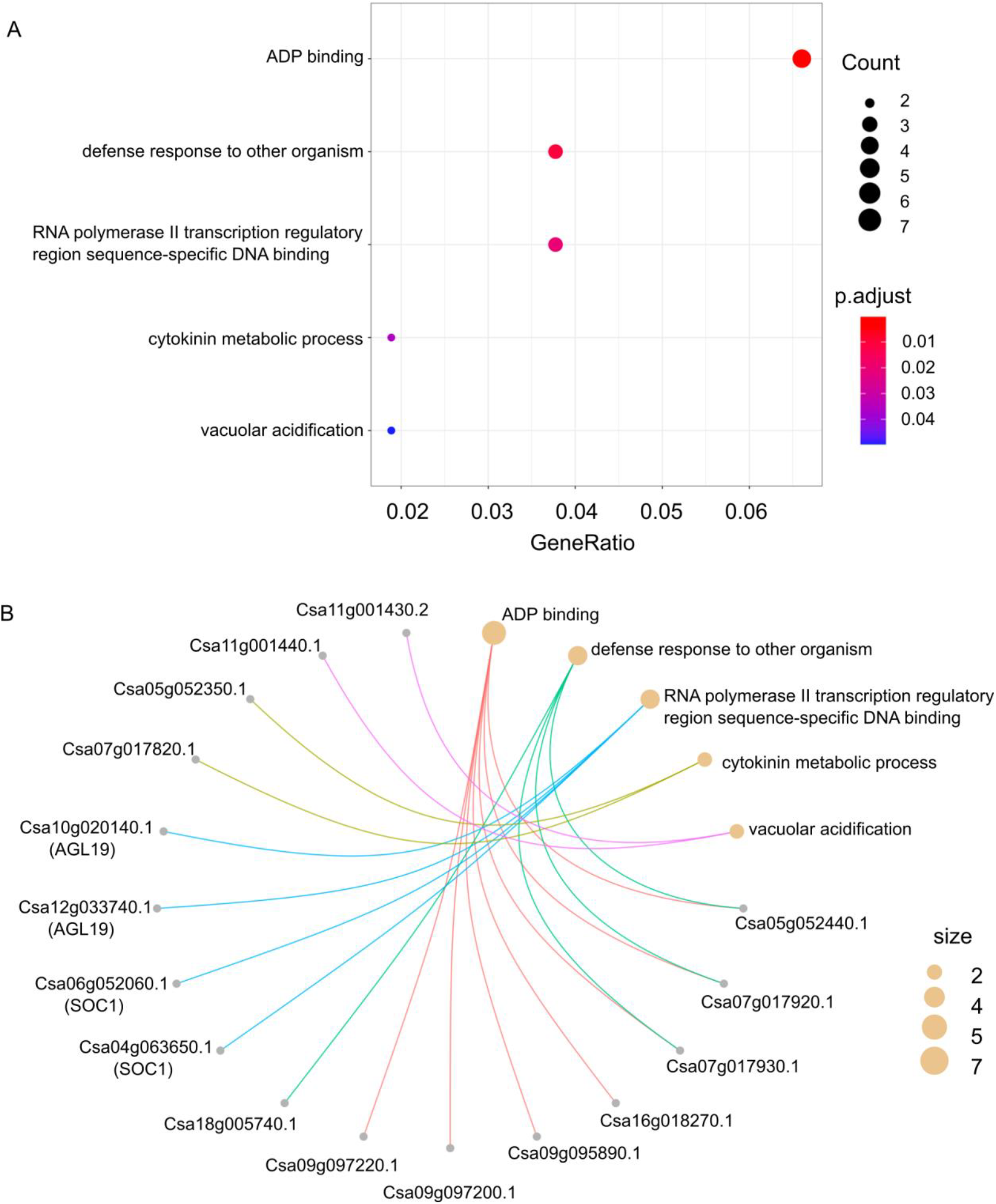
GO enrichment analysis of tan module. (A) Dot plot showing the 5 significantly enriched GO terms of tan module. Color bar indicates the p-values from Fisher’s exact test, and dots size is the number of genes belonging to the given GO term. (B) Cnetplot listing details about what *C*.*sativa* genes associated with each GO term.

### 5 eQTL and association analysis of tan module

As all 164 tan genes can be considered functionally FT-relevant genes given their high correlation with the FT trait, an expression quantitative trait loci (eQTL) analysis was performed to identify putative eQTLs that have *cis* or *trans* regulatory effects on the expression of these FT-related genes. During the eQTL analysis, all 65,082 SNPs were tested for their association with each of the 72,713 expression traits (i.e. gene expressed in least at one genotype). Accordingly, the Bonferroni adjusted p-value threshold was calculated as 0.05/ (65,082 × 72,713) = 1.06 × 10^−11^. At this threshold, in total 1,769 SNPs (eQTLs) were identified as significantly associated with genes in the tan module.

To further characterize the set of 1,769 SNPs (eQTLs) above, transcriptome-wide association studies (TWAS) were used to mine the suggestive loci that were significantly associated with the FT trait. Employing a Linear Mixed Module in GEMMA, 10 out of the 1,769 SNPs were confirmed to be significantly associated with FT (P-value < 5 × 10^−7^ ; **Figure 6A**). See **Table 2** for a comprehensive view of these 10 significant FT-associated SNPs, their related gene, and corresponding annotations. Interestingly, when we examined these 10 SNPs, their distribution across the 48 experimental population show 3 distinct patterns, forming distributed haplotypes (**Supplementary data 3**). To further check their allelic effects on FT trait, each individual haplotype was plotted against FT, which suggested that these 10 SNPs might directly contribute to the FT trait variation of *C. sativa* (**Figure 6B**). To conclude, by combining WGCNA, eQTL, and TWAS analysis, we identified 10 important candidate SNPs involved in FT variation in *C. sativa* plants.

**Table 2:** Significant SNPs associated with flowering time.

| SNP | Chr. | Pos. | Adjusted P-value | Annotation | Associated gene | Encoded protein |
| --- | --- | --- | --- | --- | --- | --- |
| snp_Ch6_18702229 | 6 | 18702229 | 1.42E-08 | missense_variant | Csa06g036830 | aquaporin NIP2.1 |
| snp_Ch6_21176774 | 6 | 21176774 | 1.42E-08 | missense_variant | Csa06g042890 | jumonji/zinc-finger-class transcription factor protein |
| snp_Ch8_24560438 | 8 | 24560438 | 1.42E-08 | synonymous_variant | Csa08g057320 | Encodes sorting nexin SNX2b |
| snp_Ch9_30922977 | 9 | 30922977 | 1.42E-08 | synonymous_variant | Csa09g081420 | 12-oxophytodienoic acid reductase |
| snp_Ch11_18992361 | 11 | 18992361 | 1.42E-08 | synonymous_variant | Csa11g040110 | phytoene desaturase |
| snp_Ch11_47854078 | 11 | 47854078 | 1.42E-08 | synonymous_variant | Csa11g101310 | Arabidopsis ortholog of human splicing factor SC35 |
| snp_Ch14_2073315 | 14 | 2073315 | 1.42E-08 | synonymous_variant | Csa14g006410 | Peptidase M50 family protein |
| snp_Ch20_8623095 | 20 | 8623095 | 1.42E-08 | synonymous_variant | Csa20g025800 | Phosphate translocator family member |
| snp_Ch20_12463032 | 20 | 12463032 | 1.42E-08 | missense_variant | Csa20g036360 | EMBRYO DEFECTIVE 1211; a MORN motif containing protein |
| snp_Ch20_12464248 | 20 | 12464248 | 1.42E-08 | missense_variant | Csa20g036360 | EMBRYO DEFECTIVE 1211; a MORN motif containing protein |

**Figure 6.**
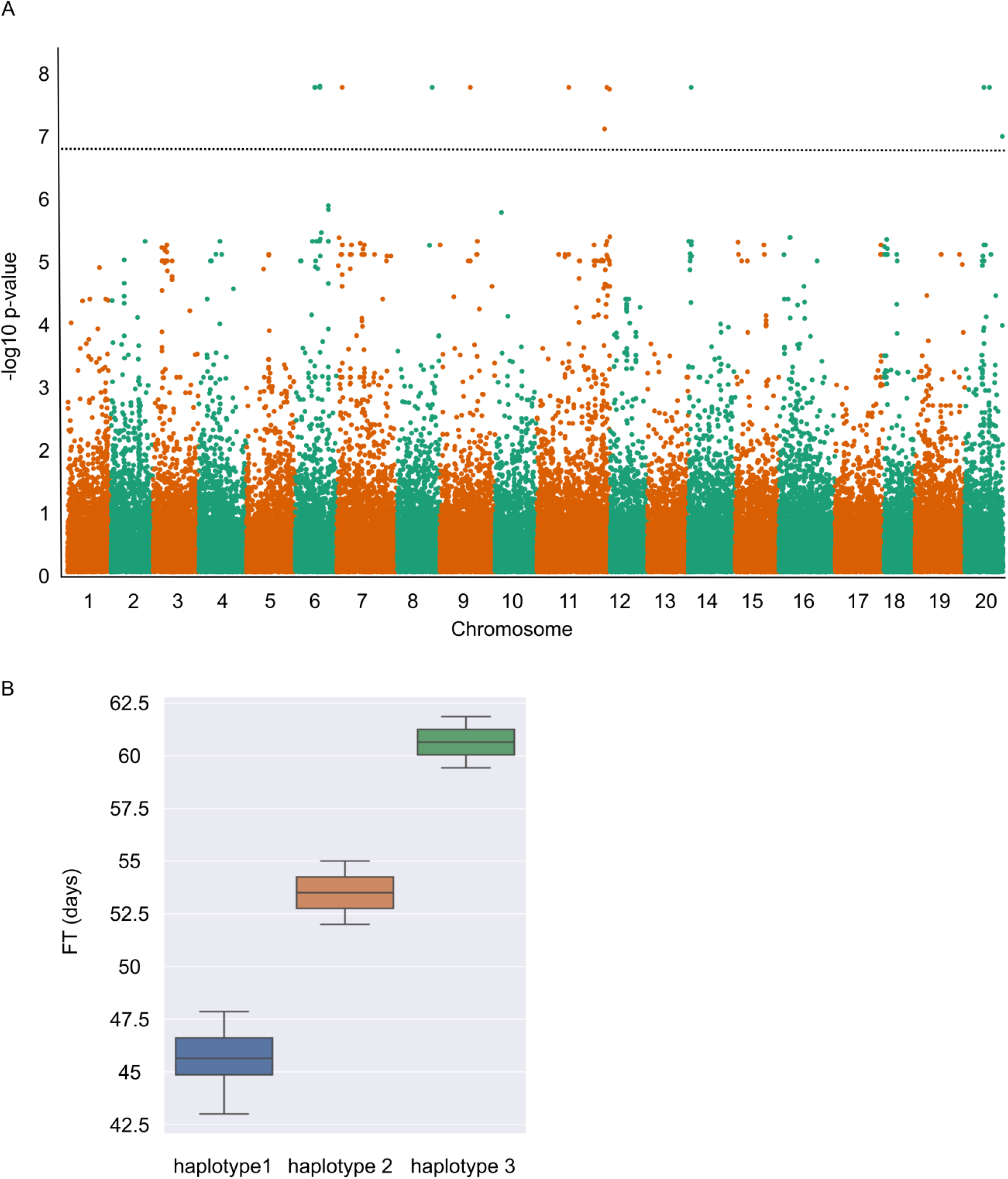
Association analysis identifies FT-related SNPs. (A) Manhattan plot showing results from association analysis, The y axis represents –log(P values) of association between each SNP variant with FT trait, and the horizontal dotted line represents the threshold for significance. (B) Boxplot showing the FT values of the *C. sativa* accessions represented by each haplotype.

## Discussion

Multiple analyses have shown *C. sativa* to be limited in genetic diversity (Chaudhary et al., 2020; Gehringer et al., 2006; Singh et al., 2015); yet there has been documented evidence of phenotypic variability in available germplasm for a number of traits (Berti et al., 2016; Luo et al., 2019). Some of the latter variation could be explained by simple single gene control; however, it is probable that the multiple layers of genetic and epigenetic control deployed by most species are being exploited in *C. sativa* to allow further adaptation. In order to gain insights into the role of the transcriptome in providing opportunities for trait adaptation, gene expression was assessed in young leaf tissue across 48 accessions of *C. sativa*. These data showed distinct patterns of gene expression among the lines; although 37,268 core genes were expressed in all lines, between 8,374 and 18,314 additional genes were variably expressed across the experimental population. Notably the most specifically expressed genes, found in ≤10 lines, were highly enriched for genes involved in plant defense; interestingly this observation aligns with that for plant pan-genomes, where genes related to biotic and abiotic stress responses are commonly over-represented among the annotated variable genes (Bayer et al., 2020). Although without full genome assemblies, we cannot rule out copy number variation (CNV) among the accessions, the transcriptome data suggests that CNV alone does not control trait variation in *C. sativa*. The polyploid genome structure of *C. sativa* also provides opportunity for trait adaptation; *C. sativa* has a highly undifferentiated hexaploid genome with most functional genes being represented by three orthologues (Kagale et al., 2014), providing opportunities for neofunctionalization of the duplicate gene copies. Among the transcriptome data 12,440 triads were identified and assessed for balanced or biased expression across the three sub-genomes. Approximately one third (37%) of the triads showed conserved expression, either balanced or biased, across all of the lines, and as noted previously for *C. sativa* the third genome showed the highest level of genome dominance in all 48 accessions. The remaining triads showed evidence of differential bias among the accessions, providing opportunities for adaptation (Wang et al., 2022). Such variation has been shown to be of importance for crop improvement in polyploid wheat, with selection for particular yield traits resulting in genome imbalance associated with low-expressing alleles of specific homoeologues (He et al., 2022). The prevalence of transcription factors among the triads showing the most variable patterns of bias among the accessions in *C. sativa* would suggest genome dominance could be a useful source of variation in this species. The cumulative transcriptome dataset provides a novel resource for studying trait variation in *C. sativa*, which was further queried to look for novel variation associated with FT in *C. sativa*.

### Floral genes in *C. sativa*

The floral integrator genes (e.g. *FLC* and *SOC1*) display very high conservation across flowering plants (Kim, 2020; Voogd et al., 2015). Further, the high levels of sequence homology and gene collinearity between *C. sativa* and Arabidopsis suggests *C. sativa* genes are expected to function in a similar manner as their Arabidopsis orthologous genes (Berti et al., 2016; Kagale et al., 2014). Study of a module highly correlated with FT from the co-expression network analysis captured two of the three *C. sativa SOC1* orthologues, but not the third. This might suggest two *Csa*.*SOC1*s work in a concerted manner during FT regulation while the third *Csa*.*SOC1* has undergone neofunctionalization, as was suggested for *Csa*.*FLC* (Anderson et al., 2018). However, study of additional *Camelina* lines has identified a functional third copy of *Csa*.*FLC*, indicating that there may be variation yet to be discovered (Chaudhary et al., 2023). Interestingly, *SOC1*-like genes (*AGL19*) also contributed two copies to the same co-expressed gene module. In Arabidopsis, *AGL14, AGL19, SOC1* and *AGL42* belong to the *TM3*/*SOC1* clade based on phylogenetic analysis (Becker & Theissen, 2003). Not only do they share high sequence similarity, intensive interactions have also been shown between these *AGL*s, e.g. *SOC1* has been demonstrated to directly regulate the expression of *AGL42* during floral transition by balancing its expression level (Dorca-Fornell et al., 2011). Further, *AGL42* activates the expression of all *AGL14/19/SOC1* genes during flower development, and all *AGL14/42/SOC1* genes show upregulated expression in the flowers of plants expressing 35S::AGL19, suggesting potential positive feedback loops between these *SOC1* and *SOC1*-like genes, such that they reciprocally activate each other in order to regulate flowering time (Lee & Lee, 2010). However, there is likely to be a more complicated regulation landscape in *C. sativa* given that *C. sativa* possesses three copies of each of the *SOC1* and *SOC1*-like genes (Kagale et al., 2014), adding another layer of complexity to the FT regulation network; how these *Csa*.*SOC1*s and *SOC1*-like genes orchestrate to fine-tune the FT of *C. sativa* will require further analyses.

### FT-associated SNPs

The 10 FT-associated SNPs could be resolved into three distributed haplotypes, one of which was dominant among the lines, while two lines carried haplotype 2 and an additional two lines haplotype 3; with number of days to flower increasing for those lines carrying the latter two haplotypes (**Figure 6B**). Since *FLC* is known to inhibit flowering by repressing *SOC1*, and its role in conferring the winter habit in *C. sativa* has already been suggested (Chaudhary et al, 2023), the lines were assessed for expression at any of the three *FLC* loci. Interestingly, all lines showed a low level of expression from the *Csa*.*FLC*.*C08* locus, and no significant expression from the C13 and C20 loci, with the exception of the haplotype 3 lines which showed significantly higher levels of *Csa*.*FLC*.*C20*, and the haplotype 2 lines which showed significantly higher levels of *Csa*.*FLC*.*C13*. It appears that there is a natural variation for expression at the FLC loci which may contribute to later flowering.

Based on the annotations of the identified 10 FT-associated SNPs, there were two SNPs (snp_Chr20_12463032 and snp_Chr20_12464248) located to the same gene (Csa20g036360), suggesting a candidate haplotype block spanning at minimum the 1216 base pairs across this gene. Further BLASTP searches showed the orthologous gene of Csa20g036360 in Arabidopsis is AT5G22640, encoding EMBRYO DEFECTIVE 1211 (EMB 1211). Previous studies have shown that EMB 1211 is a key player involved in embryo development as well as chloroplast biogenesis (Liang et al., 2010; Tzafrir et al., 2004). Notably, constitutive expression of a maize *SOC1* (*ZmSOC1*) in soybean resulted in *EMB 1211* being highly upregulated, which suggested a possible feedback loop between SOC1 and EMB 1211 during plant development (Han et al., 2021). Interestingly, the correlation between the expression of Csa20g036360 and FT is very low (r = -0.12; p-value = 0.17), which suggests its involvement in FT regulation is likely non-linear in *C. sativa*.

Several other SNPs identified in this study merit further investigation to gain a more comprehensive understanding of their roles in the regulation of flowering time in *C. sativa*. One of the significant SNPs, snp_Chr6_21176774 was located within the region of gene Csa06g042890, whose ortholog in Arabidopsis encodes a Jumonji/zinc-finger-class transcription factor protein, namely JMJ19, which contains both Jumonji N (JmjN) and Jumonji C (JmjC) domains (Lu et al., 2008; Noh et al., 2004). Based on sequence similarities, there are a total of 9 putative JmjN/JmjC-containing proteins in Arabidopsis, which are JMJ11, JMJ12, JMJ13, JMJ14, JMJ15, JMJ16, JMJ17, JMJ18, and JMJ19. These 9 JmjN/JmjC-containing proteins are further divided into 2 sub-groups, whereat JMJ11, JMJ12 and JMJ13 belong to the KDM4/JHDM3 group and the other six belong to the KDM5/JARID1 group (Lu et al., 2008). JMJ11 and JMJ12 are known as EARLY FLOWERING 6 (ELF6) and RELATIVE OF EARLY FLOWERING 6 (REF6) respectively, which have been reported to play divergent roles in flowering regulation in Arabidopsis (Noh et al., 2004); as their homolog, JMJ13 has also been shown to be a flowering repressor dependent on temperature and photoperiod (Zheng et al., 2019). Furthermore, JMJ14 acts through repression of floral integrators (i.e. FT, AP1, SOC1 and LFY) to prevent early flowering during the vegetative phase (Jeong et al., 2009; Lu et al., 2010). As a large number of histone demethylases contain a conserved Jumonji C (JmjC) domain, these 9 proteins were predicted to be potential histone demethylases, and function in epigenetic regulation (Jeong et al., 2009; Lu et al., 2010; Lu et al., 2008; Zheng et al., 2019), which was affirmed by (Noh et al., 2004) where they showed JMJ12 functions as an FLC repressor through histone modifications of FLC chromatin. Therefore, it is possible that JMJ19 functions in a similar manner, as a histone demethylase during the regulation of flowering in *C. sativa*. Interestingly, snp_Chr11_47854078, locates to a region harboring the ortholog of Arabidopsis *HUMAN SPLICING FACTOR SC35* (*AtSC35*). AtSC35 is a member of the serine/arginine-rich (SR) proteins, which are important splicing factors (Barta et al., 2010). AtSC35 has been shown to play role in pre-mRNA splicing along with other four SC35-like (SCL) proteins. Further, AtSC35 and SCL proteins, in a redundant manner, control the transcription as well as the alternative splicing of *FLC* to regulate flowering time in Arabidopsis (Wang et al., 2023; Yan et al., 2017).

### Conclusion

FT is an important agricultural trait and understanding the mechanisms underlying its control could offer practical solutions in adapting FT to improve crop productivity in different environments. Owing to the identified allohexaploid genome, the genetic and molecular basis of FT are only beginning to be uncovered in *C. sativa*. In this study, WGCNA, eQTL and TWAS analysis were combined to dissect the FT trait in *C. sativa*; the results provide not only new insights into the genetic basis of *C. sativa* FT trait, but also offer new target genes for genetic improvement during future *C. sativa* breeding. Interestingly, the FT-associated SNPs identified do not overlap those from a previous study (Luo Lily et al., 2021) but did show some correspondence to the expression of *FLC*. As our SNPs result from expression data, while (Luo Lily et al., 2021) carried out GWAS based on SNPs from genotyping-by-sequencing (GBS), this might explain the lack of correspondence. These results also indicate the complex genetic architecture underlying FT regulation in the allohexaploid plant *C. sativa*, and emphasize the necessity of employing additional approaches to help us verify the core elements required for floral transition in *C. sativa*.

## Materials and methods

### Plant materials and phenotypic measurements

A *C. sativa* population comprising of 48 different accessions collected by Plant Gene Resources of Canada (PGRC, Saskatoon, SK, Canada) were used in this study. All accessions were planted in field trials at experimental farm on Lowe Road, Saskatoon, during the growing season of 2012 and 2013. Three replicates were grown for each accession per season, double rowed plots. Flowering time (FT) was measured when 50% of plants had flowered in each plot. Inflorescence length, internode length and plant height were recorded as the average of three measurements per plot. Mature pods and seeds were collected from three plants in each plot to measure pod shattering and 1000-seed weight respectively.

### RNA extraction and sequencing

The fully expanded true leaves from the young seedlings of *C. sativa* were collected to extract total RNA using the RNeasy plant mini kit (Qiagen) according to the manufacturer’s protocol. The concentration and integrity of total RNA were examined by BioAnalyzer with RNA 6000 Nano Kit (Agilent). cDNA sequencing libraries were constructed according to the standard TruSeq RNA library preparation guide, and paired-end sequencing was performed using the HiSeq 2000 platform (Illumina).

### Expression profiling and SNPs calling

Raw RNA-seq data were filtered using Trimmomatic (version 0.32)(Bolger et al., 2014) with the following settings (ILLUMINACLIP:TruSeq3-PE.fa:2:30:10; SLIDINGWINDOW:4:15; LEADING:15; TRAILING:15; MINLEN:55) to remove adapter and low-quality sequences. For expression profiling, clean reads were aligned to the *C. sativa* reference genome using STAR (version 2.7.10a)(Dobin et al., 2013) and gene expression levels were summarized by featureCounts (version 2.0.1)(Liao et al., 2014). Subsequent raw counts were filtered to remove lowly-expressed genes (less than 1 across the samples), and then normalized through variance stabilizing transformation (VST) in DESeq2 (Love et al., 2014). The number of expressed genes across all 48 genotypes were calculated by assigning a gene as expressed only if the normalized counts were greater than 3 in all three biological replicates.

For SNP calling, STAR’s two-pass mode was employed to map clean RNA reads to *C. sativa* reference genome. The binary alignment map (BAM) files were used to make unmapped BAM (uBAM) files through Picard’s RevertSam function (version 2.27.4), filtered to remove PCR duplicates through Picard’s MarkDuplicates function and further processed by the SplitNCigarReads function in GATK (version 4.2.6.1)(McKenna et al., 2010). The retained reads were used for variant calling through GATK’s HaplotypeCaller, CombineGVCFs, and GenotypeGVCFs functions with default parameters. Next, SNPs were collected using GATK’s SelectVariants function, and filtered through the VariantFiltration function with setting “QD < 2.0 || MQ < 40.0 || FS > 60.0 || SOR > 3.0 || MQRankSum < -12.5 || ReadPosRankSum < -8.0”. Eventually, all 1,161,502 SNPs were further filtered through vcftools (version 0.1.16)(Danecek et al., 2011) with parameters “max-missing 0.95, maf 0.05, hwe 0.000001”, and 65,082 SNPs were obtained for further analysis.

### Dominant Expression/Genome Bias

To identify the highly variable genes that are significantly dominantly (or specifically) expressed in select lines, TAU index was used for defining a gene’s specific expression score. TAU index of each gene from top expressed n lines (1≤n≤10) were calculated respectively and TAU>0.9 was used as a threshold to define the dominantly expressed genes in a few lines. The highly variable genes identified based on TAU index were further visualized using hierarchical clustering combined with a gene expression heatmap performed by R pheatmap package (https://cran.r-project.org/web/packages/pheatmap/index.html).

Normalized counts of 12,440 syntenic triads from all RNA-seq samples were used to study the orthologous expression bias in all 48 *C. sativa* accessions based on the definition in a previous study (Ramírez-González et al., 2018). With the thresholds defined as 20, 80, and 100%, seven patterns for expression bias, including “Balanced”, “A dominant”, “B dominant”, “C dominant”, “A suppressed”, “B suppressed”, “C suppressed”, were identified based on the expression difference among the syntenic triads from the three sub-genomes. All patterns were distinguished for the triads with at least one member holding an average expression of 10 counts across all 48 *C. sativa* accessions. Ternary plots were used to visualize the relative expression contribution of each member of triads from three sub-genomes and plotted by R ggtern package (Hamilton & Ferry, 2018).

### WGCNA and GO enrichment analysis

The VST normalized counts were further used to extract a subset (15,000) of the most highly variable genes across the experimental population through genefilter package in R. The selected subset of 15,000 most variable genes were used to construct a co-expression network through the WGCNA package in R (Langfelder & Horvath, 2008), whereat the function blockwiseModules was used with parameters “power = 12, maxBlockSize = 15000, networkType = “signed”, TOMType = “signed”, minModuleSize = 30, mergeCutHeight = 0.25” to construct a signed network and generate co-expressed gene modules. The *C. sativa* GO annotation database was created with its whole proteome using InterProScan 5 (Jones et al., 2014). Further, GO enrichment analysis was conducted for the module of interest using clusterProfiler in R (Wu et al., 2021).

### eQTL and association analysis

eQTL analysis was performed through Matrix eQTL in R (Shabalin, 2012). 65,082 SNPs and expression data from the selected subset of 15,000 genes were used to perform the eQTL analysis. Significant eQTLs were defined as *cis* when the physical distance between SNP and its associated gene was less than 3 kb, and *trans* if the distance exceeded 3 kb. For transcriptome-wide association study (TWAS), a mixed linear model through program GEMMA (version 0.98.5)(Zhou & Stephens, 2012) was carried out to detect the associations between 65,082 SNPs and FT’s Best Linear Unbiased Predictors (BLUPs) values, which were generated using lme4 (version 1.1-34) in R. The first three principal components derived from all 65,082 SNPs were used as a covariate in the model to correct for population stratification.

## Data Availability

The raw RNA-seq data have been deposited to the Gene Expression Omnibus (GEO) under the accession GSE253393. All other data are included in the article and/or supporting data.

## Acknowledgements

This work was supported through funding provided by Saskatchewan Agricultural Development Fund, and by funding to Genome Prairie from the Western Economic Partnership Agreement project ‘Prairie Gold’. The authors would like to thank Christina Eynck and her team at Saskatoon Research and Development Centre for their assistance in the field.

## Figures

**Figure S1.**
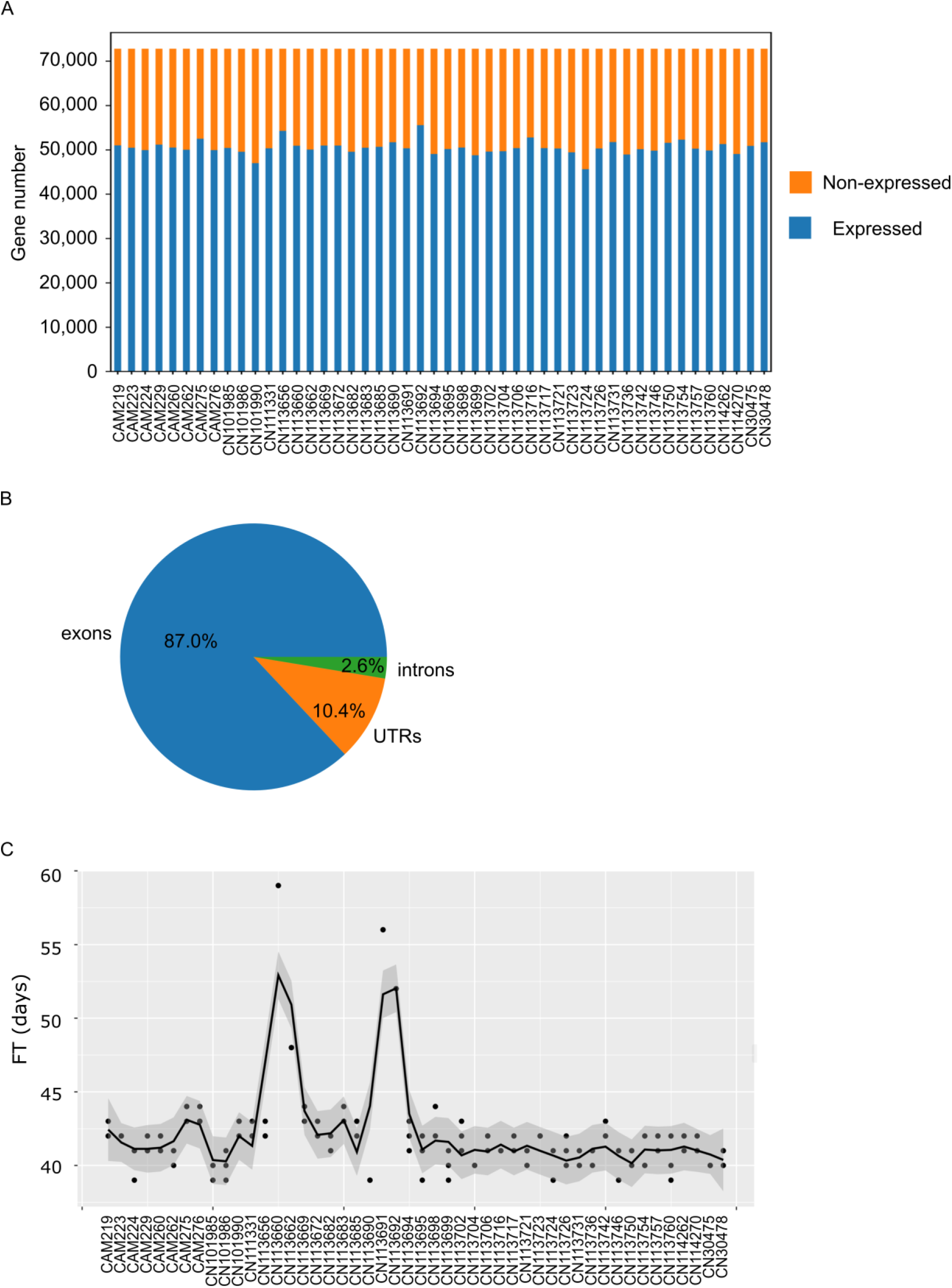
Gene expression and FT across various genotypes and SNPs categories. (A) gene expression variation across 48 *C*.*sativa* accessions. For each genotype, expressed gene was defined as its average expression of 3 replicates ≥ 5 vst normalized counts. (B) Pie plot shows the percentage of all 65,082 SNPs belonging to each category. (C) Smoothed line plot showing the FT variation across all 48 genotypes in year 2013.

**Figure S2.**
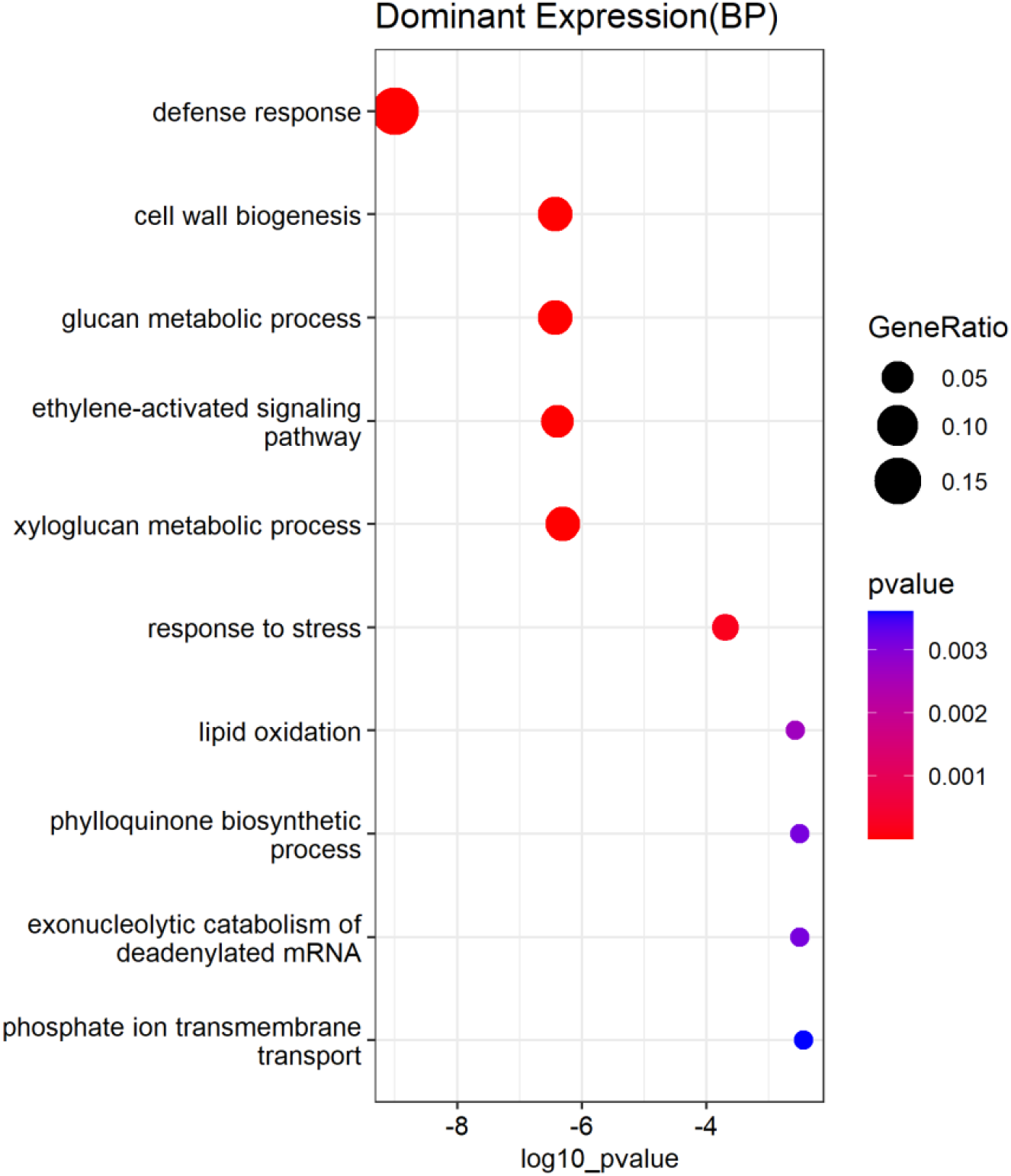
Enrichment for Biological Processes among Gene Ontology Annotation of most dominantly/specifically expressed genes.

